# A molecular framework for control of oriented cell division in the Arabidopsis embryo

**DOI:** 10.1101/2021.02.09.430440

**Authors:** Prasad Vaddepalli, Thijs de Zeeuw, Sören Strauss, Katharina Bürstenbinder, Che-Yang Liao, Richard Smith, Dolf Weijers

**Affiliations:** Laboratory of Biochemistry, Wageningen University, Stippeneng 4, 6708WE Wageningen, the Netherlands; Department of Comparative Development and Genetics, Max Planck Institute for Plant Breeding Research, Carl-von-Linne-Weg 10, Cologne, Germany; Department of Molecular Signal Processing, Leibniz Institute of Plant Biochemistry, Weinberg 3, Halle (Saale), Germany; John Innes Centre, Norwich Research Park, Norwich, United Kingdom; Laboratory of Plant Physiology, Wageningen University, Droevendaalsesteeg 1, Wageningen, the Netherlands; Plant Ecophysiology, Institute of Environmental Biology, Utrecht University, Utrecht, the Netherlands

## Abstract

Premitotic control of cell division orientation is critical for plant development, as cell walls prevent extensive cell remodelling or migration. Whilst many divisions are proliferative and add cells to existing tissues, some divisions are formative, and generate new tissue layers or growth axes. Such formative divisions are often asymmetric in nature, producing daughters with different fates. We have previously shown that in the *Arabidopsis thaliana* embryo, developmental asymmetry is correlated with geometric asymmetry, creating daughter cells of unequal volume. Such divisions are generated by division planes that deviate from a default “minimal surface area” rule. Inhibition of auxin response leads to reversal to this default, yet the mechanisms underlying division plane choice in the embryo have been unclear. Here we show that auxin-dependent division plane control involves alterations in cell geometry, but not in cell polarity or nuclear position. Through transcriptome profiling, we find that auxin regulates genes controlling cell wall and cytoskeleton properties. We confirm the involvement of microtubule (MT)-binding proteins in embryo division control. Topology of both MT and Actin cytoskeleton depend on auxin response, and genetically controlled MT or Actin depolymerization in embryos leads to disruption of asymmetric divisions, including reversion to the default. Our work shows how auxin-dependent control of MT- and Actin cytoskeleton properties interacts with cell geometry to generate asymmetric divisions during the earliest steps in plant development.

## Introduction

Plants rely heavily on precise control of cell expansion and division plane selection since their cells are encased by rigid cell wall and cannot migrate from their position. Mechanisms controlling the division plane orientation have been an area of focus for over a century [1–3]. Starting from the establishment of the early embryo to the development of post-embryonic tissues and organs from meristematic tissues, plants need to constantly calibrate the co-ordination between cellular and genetic inputs for proper cell and tissue patterning. Failure in the co-ordination leads to aberrant phenotypes with severe developmental defects [4–6]. Proliferative mitotic cell divisions select symmetric division plane resulting in cells with approximately equal size. In formative divisions, however, division planes strongly deviate from the symmetric position, leading to daughter cells of different sizes. Such asymmetric divisions often lead to the formation of new cell identities and tissue layers, and these divisions can thus lead to differential developmental fate in addition to unequal volume partitioning.

In plants, the cortical microtubule (CMT) array is linked with the direction of division plane orientation [7]. CMTs condense into a thick band and, together with Actin, form a plant-specific cytoskeletal structure called the preprophase band (PPB), forecasting the future division plane [8]. PPB formation involves changes in cytoskeletal dynamics and stabilization. Several cytoskeleton-associated and regulatory protein complexes involved in this process have been identified [9]. Although the relevance of the PPB for controlling division plane is debated, it is clear that the structure is a good predictor of division plane, and required for robustness [10], which is crucial for proper asymmetric divisions. So far, the mechanisms that connect developmental regulators with CMTs and actin to influence the positioning of the CMT and PPB are largely unknown. How the cytoskeleton integrates cell geometry and other regulatory input into division plane orientation remains a mostly unanswered question.

Plant cells by default divide along the minimal surface area (in 3D) following the “shortest-wall” (in 2D) rule [11]. Cell geometry therefore is a fundamental input in deciding the size and shape of the daughter cells. Genetic elements interfere with the default symmetric division to facilitate division plane orientation [12]. Recent evidence implicate that cytoskeleton dynamics may bridge the co-ordination of geometric and genetic input to influence the re-orientation of the division plane [13,14]. During the first asymmetric division of the zygote and in lateral root founder cells, dynamics of cytoskeletal pattern decide the correct orientation of division plane. In both these systems however, cells are elongated, and the various orientations of division are dramatically different in terms of surface area and volume partitioning. A key question is whether similar mechanisms operate in the smaller, polyhedral cells where such differences are less extreme. The signalling cue for biasing division plane orientation likely involves cell polarity mechanisms [15], but how the intracellular position of the polarity proteins direct division plane orientations remains elusive. In several cell types, the nuclear position co-aligns with the PPB [16], and migration of the nucleus is correlated with positioning of the division plane wall in the zygote, lateral root founder cells and in leaf epidermis [13,14,17]. Again, all those cell types are either large, relative to nuclear size, or have extreme aspect ratios, and it remains a question whether the same principles apply to division control in other types of cells.

Developing from a fertilized egg cell, the early plant embryo is a hotspot for formative events: new cell types are established with many divisions, which in Arabidopsis are highly predictable [18,19]. With the advent of advanced imaging and cellular segmentation approaches, a 3D description of early Arabidopsis embryogenesis has been generated [12]. From this work, it surfaced that divisions leading to the 2, 4 and 8 cell embryo stages follow the minimal surface area rule, corroborating classical (2D) models from 19th century. However, the next, asymmetric divisions that generate the protoderm and inner cells at the 16-cell stage deviated from this rule. Using mutant embryos in which the response to the plant hormone was blocked by ubiquitous expression of a transcriptional repressor (*RPS5A>>bdl;* [20]), it was demonstrated that transcriptional response to the auxin hormone is required to suppress the geometric default division, implicating that the regulation of oriented cell division by geometric and genetic cues can be uncoupled. Thus, the 8-cell Arabidopsis embryo represents a unique case where the activity of a transcriptional regulator (bdl) allows to switch between default, symmetric and regulated asymmetric division. Based on a more recent computational model, it has been proposed that all division planes observed in wild-type and mutant cells conform to a default rule, provided that new walls can be curved when inserted [21]. In the same study, it was suggested that also in these cells, nuclear position may provide input into division plane choice [21]. Analysis of live embryos found little to no curvature in newly formed walls [12], and it is therefore an entirely open question through what cellular processes, genes and mechanisms division orientation in early embryos is controlled. Here, we explore mechanisms underlying division plane selection in the embryo. Using computational approaches and genetic perturbation, we studied the role of cell polarity, nuclear position and cell shape in determining the division plane orientation. In addition, we identified a set of auxin-dependent genes involved in division plane orientation, which revealed cell wall and cytoskeleton regulating genes. We investigate the potential role of IQD6 protein clade in division plane orientation. Further, we show that cytoskeleton dynamics are critical contributors to auxin-dependent early embryo division plane orientation.

## Results

### TIRl/AFB-dependent auxin response controls cell division orientation in the early embryo

In the nuclear auxin pathway, presence of auxin leads to the degradation of AUX/IAA proteins by TIR1/AFB receptors and thus promoting ARF-dependent gene expression [22]. A mutation in the degron of Aux/IAA proteins prevents auxin-dependent interaction with TIR1/AFB proteins, and causes accumulation of the mutant protein, thus leading to permanent inhibition of ARF proteins [20,23]. We have previously shown that broad expression of the auxin-insensitive mutant iaa12/bdl protein in early embryos (*RPS5A>>bdl*) prevents asymmetric divisions at the 8-cell stage [12]. However, since the mutant bdl protein can accumulate to unnaturally high levels, this may lead to inhibition effects beyond the normal activity of auxin. It is therefore not clear if an endogenous auxin response process controls division orientation. To address this question, we scrutinized 3D division orientation in mutant embryos lacking all 6 TIRl/AFB receptors, the *tir1/afb* sextuple (*tir1afb12345*) mutant [24]. Since this mutant was shown to arrest during embryogenesis, we made use of a sextuple homozygote that carries a heterozygous complementation transgene carrying TIR1-mOrange2::AFB5-mCherry::AFB2- mCitrine [24]. The hemizygous *tir1afb12345* mutant has 25% sextuple mutant progeny, which display division plane defects in the early embryos [24]. We performed cell segmentation and analysis using MorphoGraphX [12,25] on these mutant embryos, which revealed division plane defects closely resembling the *RPS5A>>bdl* phenotype (Fig 1A). Division plane orientation did however show variability. Next, we measured volumes of pairs of sister cells at 4-, 8-, and 16-cell stages to determine the volume distribution ratios as a proxy for division (a-)symmetry. In addition, we analysed the division plane surface area relative to the minimal and maximal planes cutting through the center of the actual division to see if divisions now approximate the 3D equivalent of the “shortest wall” [12]. In *tir1afb12345* mutant embryos, divisions leading to 4-cell and 8-cell embryos are symmetric and use the minimal surface area, similar to wild-type and *RPS5A>>bdl* embryos (Fig 1B). As reported previously, divisions leading to the 16-cell stage in *RPS5A>>bdl* embryos followed the minimum surface area giving rise to symmetric divisions, while wild-type embryos use a much larger division surface area and divide asymmetrically (Fig 1B) [12]. In *tir1afb12345* mutant embryos, divisions leading to 16-cell embryos show a high variation in normalized division plane surface area ranging between the values seen for *RPS5A>>bdl* and those in wild-type (Fig 1B). The average division is therefore more symmetric than wild-type. The results match the observed variation in division plane angles in 3D-imaging and indicate that the *tir1afb12345* mutant phenotype represents a spectrum of division plane defects that clearly include those observed in *RPS5A>>bdl* embryos, but also display weaker aberrations. It is unclear if this is due to residual TIR1/AFB activity in the hextuple mutant. Nonetheless, this analysis shows that endogenous auxin response is required for promoting asymmetric cell division in the embryo through regulating division plane orientation.

**Figure 1:**
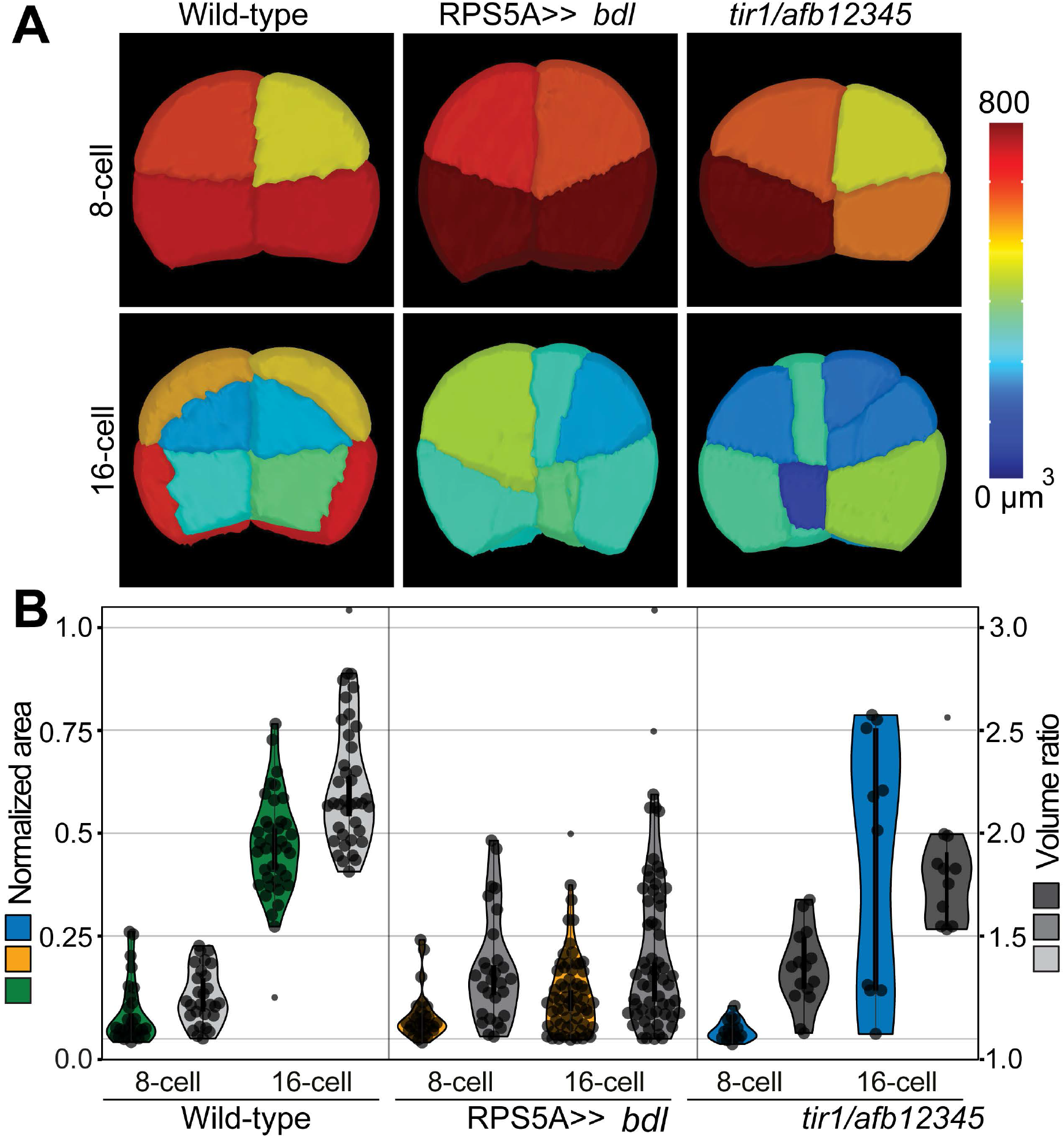
Auxin response is required for asymmetric embryonic cell division. (A) 3D comparison of wild-type, *RPS5A>>bdl* and *tir1/afb-mutant* embryos. Mesh colour per cell corresponds to cellular volume indicated in the colour scale. (B) Violin plots representing distribution of division plane areas as a fraction of the smallest (0 on the left y-axis) and largest (1 on the left y-axis) division wall area through the center of the merged volume of two sister cells. Wild-type values are shown in green, *bdl*-mutant (*RPS5A>>bdl*) values are shown in yellow, *tir1/afb* values are shown in blue. The cell volume ratios of the daughter cells resulting from these divisions are represented in grey (light to dark), and values are on the right y-axis. Individual values are shown in the violin plots. At least 3 individual embryos were used per condition.

### Cell shape, not polarity or nuclear position, correlates with division orientation

Since outer-inner cell polarity is established early in wild-type embryos [26], and because divisions at the 8-cell stage in wild-type are aligned with this polarity axis, it is conceivable that the division defects in *RPS5A>>bdl* embryos reflect a loss of polarity. We addressed this question by imaging the*pWOX2::BOR1-mCitrine* marker (ACE-W03; [26]) in *RPS5A>>bdl* and wild-type control (*RPS5A>>*Col) embryos. This marker was shown to localize to inner plasma membrane domains from the 2-cell stage onward [26], Despite characteristic defects in cell division orientation, no difference in BOR1-mCitrine localization could be detected between *RPS5A>>*Col and *RPS5A>>bdl* embryos (Fig 2A). As in wild-type embryos, the BOR1 marker is enriched at the inner cell membranes of defective 8-cell and 16-cell stage *RPS5A>>bdl* embryos. Thus, early outer-inner polarity establishment is independent of transcriptional auxin response and the failure to divide asymmetrically in the mutant is likely not caused by global loss of polarity.

**Figure 2:**
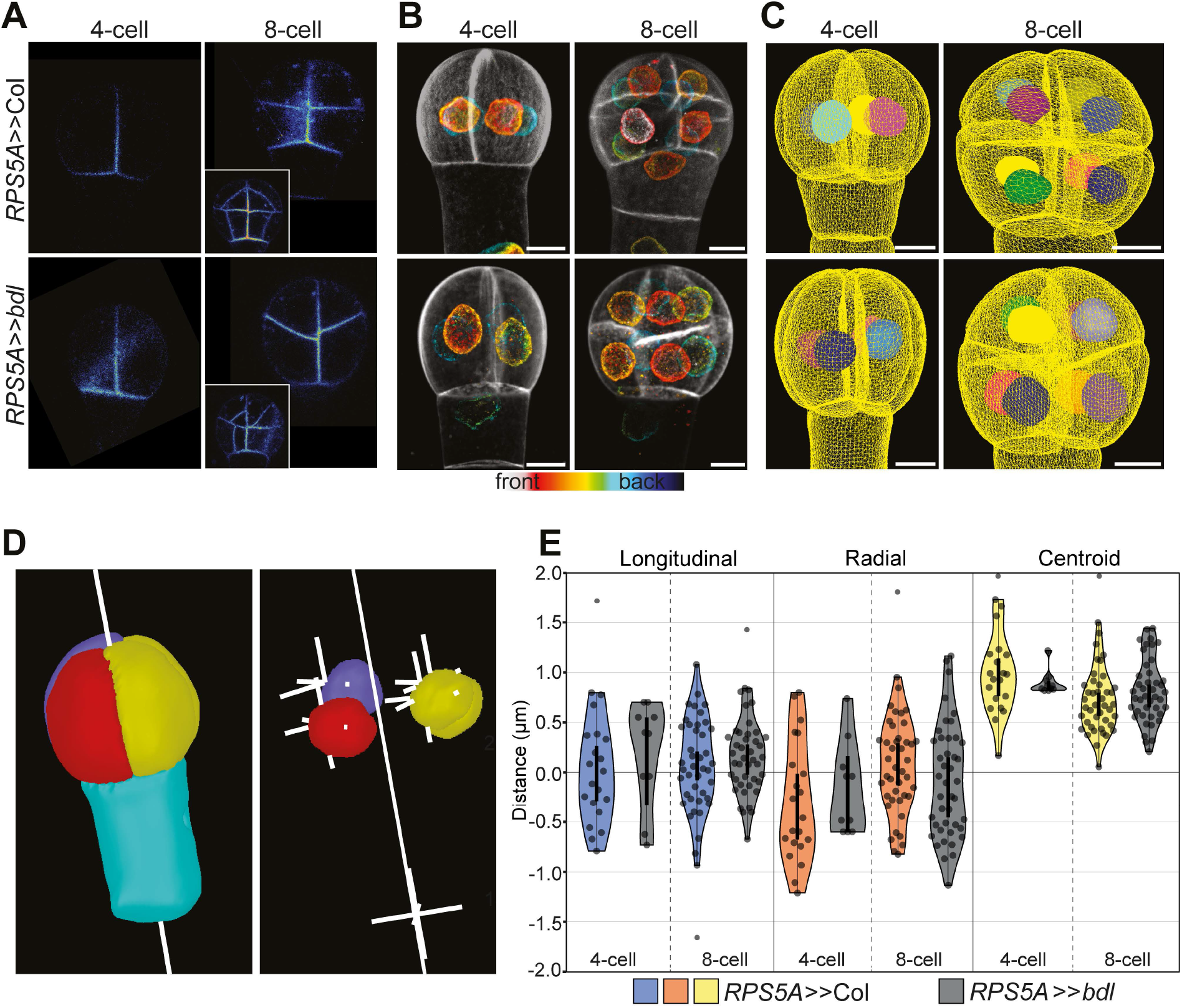
Analysis of polarity and nuclear position in wild-type and *bdl* embryos. (A) Single optical sections of embryos expressing the inner membrane marker BOR1-mCitrine (ACE-W03) in wild-type and *bdl* 4- and 8-cell embryos. Inset depicts 16-cell stage. (B-E) Nuclear position analysis in 4- and 8-cell wild-type and *RPS5A>>bdl* embryos. (B) Maximum intensity projection of depth-coded stacks of nuclear envelopes labeled by AtNUP54-GFP (ACE-W11) reporter line with SCRI Renaissance 2200 (SR2200)-stained cell walls. (C) Visualization of segmented nuclei within segmented cell meshes. Scale bars: 5 μm. (D) A central axis is defined through the embryo proper for analysis of nuclear position within the cell. From this axis, longitudinal and radial distances from each cell centroid are defined. (E) Analysis of nucleus position relative to cellular centroid position within embryos. Average distances are shown for individual cells. The distance to the centroid is absolute and not directed, and therefore cannot be below 0. Measurements were done on 9 to 43 individual cells and corresponding nuclei from at least 4 to 8 different individual embryos per condition.

Asymmetric cell division in the zygote, in lateral root founder cells and in meristemoid mother cells involve nuclear migration to the future division site, implying a strong association of division plane with nuclear position [13,14,17,27]. Compared to these systems, early embryonic cells have distinct cell geometry and our previous observations on wild-type embryos suggested that the nucleus occupies a relatively large part of the cell, limiting its ability to move compared to other larger cells [26]. However, a recent report proposed that nuclear position could constrain and guide division orientation in early Arabidopsis embryo cells [21], but given the relatively large nuclear volume, it is unclear how well its precise position correlates with division plane choice. We explored this hypothesis by determining the correlation between nuclear position and division plane in wild-type and *RPS5A>>bdl* embryo cells, where division switches between asymmetric and symmetric. We introduced the embryonic nuclear envelope marker*pWOX2::NUP54-GFP* (ACE-W11; [26]) into the *RPS5A- GAL4* background and crossed this line with wild-type or *UAS-bdl* to visualise the nuclear volume in *RPS5A>>bdl* and wild-type control embryos, High-resolution Z-stacks of early embryos were generated simultaneously for the nuclear GFP signal and for a cell wall dye. Firstly, we did not observe conspicuous differences in nuclear morphology between the two genotypes (Fig 2B). To analyse nuclear position relative to the cell volume, we created nuclear and cell outline meshes by applying MorphoGraphX-based segmentation on the Z-stacks (Fig 2C). We defined the cellular- (CC) and nuclear centroid (NC) to calculate nuclear position relative to the centroid of the cell. Defining a central axis through the embryo suspensor, we could measure general distance of NC to CC, as well as its displacement in longitudinal and radial directions (Fig 2D). For both 4-cell and 8-cell embryos, we could not find significant differences in nuclear position between wild-type and *RPS5A>>bdl-mutant* embryos (Fig 2E). Importantly, we found considerable variation in nuclear position even in wild-type cells, where the division plane is essentially invariable. These findings suggest that nuclear position may not be strongly connected to cell division orientation and is perhaps not a mechanism mediating its control.

Given that minimal surface area and the cell centroid are defined by cellular shape, a switch to a different cell division plane in auxin-insensitive mutants could also be indirectly caused by altered cell shape. Therefore, we asked if cell geometry is altered in *RPS5A>>bdl* mutant embryos before the switch to symmetric division at 8-cell stage. Interestingly, in *RPS5A>>bdl* mutants, slightly oblique division planes are observed in 8-cell stage embryos [28]. Segmentation analysis at these stages revealed that cell surface area and cell volume ratio are significantly larger in both 4-cell and 8-cell *RPS5A>>bdl-mutant* embryos compared to wild-type (Fig 3A), indicating that cell geometry is indeed affected in the *bdl*-mutant. To determine if the observed cell expansion is random or directed in either of the cellular directions, we measured circumferential-, radial-, and longitudinal cell lengths of the same embryonic cell volumes (Fig 3B). Although cell length in *RPS5A>>bdl* embryos was slightly increased in comparison to wild-type in all measured directions, most of the expansion is in the longitudinal direction in both 4-cell and 8-cell embryonic cells (Fig 3C). This result shows that the altered cell division planes are preceded by changes in cell geometry, suggesting that the primary target of auxin control could be a process that controls cell shape.

**Figure 3:**
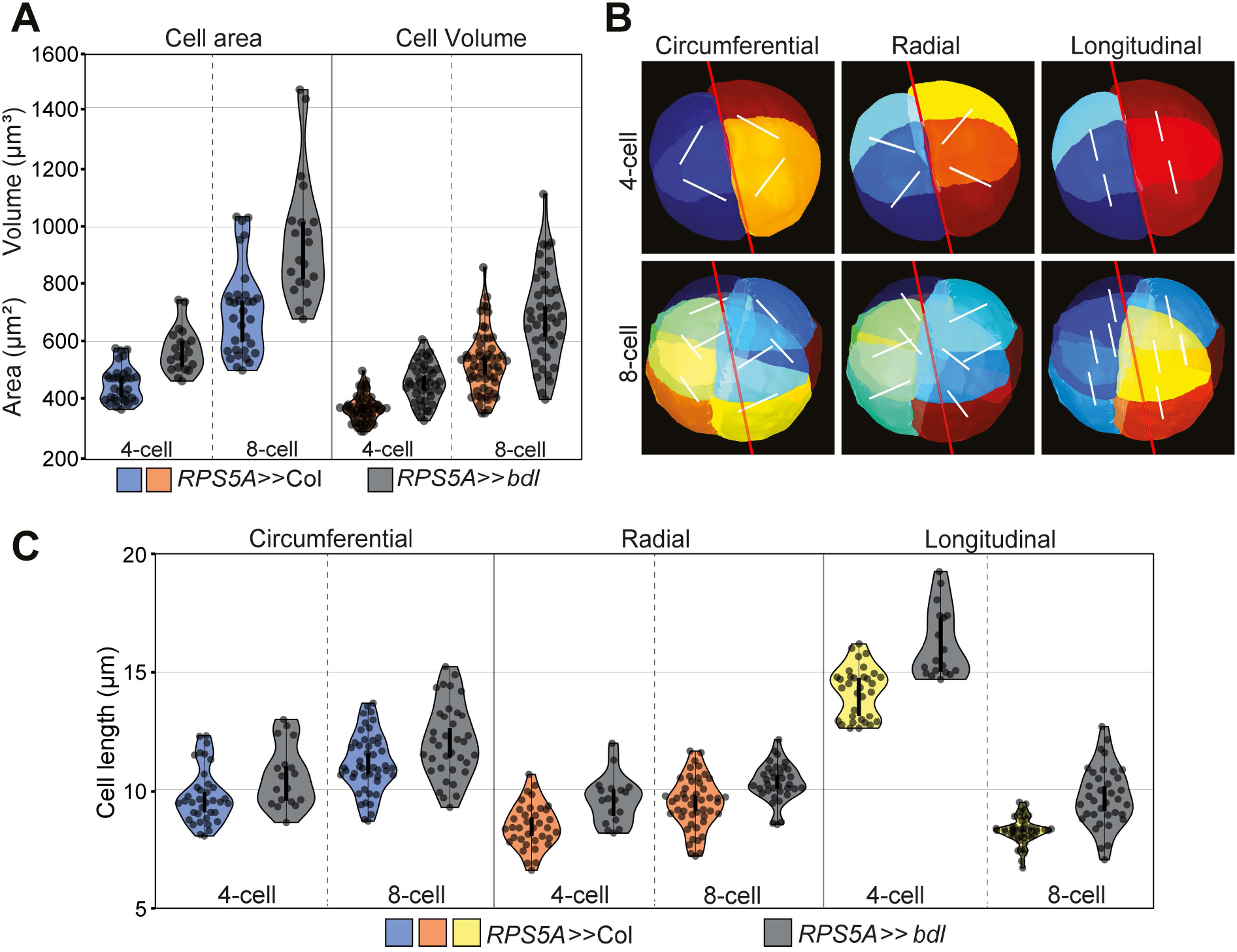
3D cell shape analysis of wild-type and *bdl* embryos. (A) Average cell surface area (in μm^2^) and cell volume (in μm^3^) are shown. Based on two-sided student t-tests, all comparisons between wild-type and mutant are statistically significant (p<0.001). Measurements were done on 20- to 56 individual embryonic cells from at least 5 individual embryos per condition. (B) Segmented wild-type embryos indicating the axis for measurements shown in (C). First, a central axis (red line) was defined for the pro-embryo. Relative from this axis, circumferential, radial, and longitudinal length measurements were performed through the cell centroid. (C) Average circumferential, radial and longitudinal cell sizes (in μm) are shown. Based on two-sided student t-tests, all comparisons are statistically significant (p<0.03). Measurements were done on 20- to 48 individual embryonic cells from at least 5 individual embryos per condition.

### Transcriptome analysis reveals altered cytoskeletal gene expression

To probe the genetic mechanisms underlying auxin-dependent cell expansion and division plane orientation, we performed transcriptome analysis, comparing manually isolated 8-cell wild-type and *RPS5A>>bdl* mutant embryos. Given that the molecular target of BDL is ARF-dependent transcriptional control, the immediate cellular pathways that are subject to auxin regulation should be apparent from the genes misregulated. We chose the 8-cell stage as this is the moment shortly before the switch in division orientation is most apparent. Initial inspection of the *RPS5A>>bdl* transcriptome revealed the expected upregulation of *BDL/IAA12* while other *Aux/IAA* genes show downregulation (Fig S1), consistent with genome-wide dampening of auxin response. Additionally, 5 out of 11 *YUC* genes were upregulated in *bdl* embryos (Fig S1), which shows that also auxin-dependent gene repression is inhibited in mutant embryos and validating the effectiveness of the inhibition of auxin response. After statistical analysis, we retained 421 up- and 414 down-regulated genes in *RPS5A>>bdl* embryos (>2-fold difference; q-value < 0.05; Supplemental Data Set 1). Gene Ontology (GO) analysis did not identify obvious enrichment of functional categories. Nevertheless, among the highly misexpressed genes, we found several genes involved in cellular mechanisms, along with known developmental regulators. Here we focus on 34 candidate genes that could be divided into three groups based on their ontology information and functional data from earlier studies (Fig 4A). The first group represents genes related to auxin signalling (*IAA1, IAA9, IAA30*), biosynthesis (*YUC1, YUC8*), and transport (*PIN1, PIN4, LAX2, NPY5*) The second group includes transcription factors, of which most are known to be key regulators of development, including several known auxin response targets (e.g. *TMO3, GATA20, WIP2, TMO5*). The third group contained genes known for their function in cytoskeletal organization and signalling, along with genes involved in cell wall composition and remodelling. A pectin methyl esterase (*PME44*), xyloglucan endotransglycosylase (*XTH19*), cellulase (*CEL2*) and an arabinogalactan protein (*FLA12*) were found downregulated in *bdl* embryos. All these are known for their roles in cell wall remodelling mechanisms during post-embryonic growth. We found significant downregulation of the ROP activating guanine exchange factor, *ROP-GEF5* along with *ROP9*, which belongs to Type II sub-group of *ROP* gene family. The plant-specific small Rho GTPase switches, ROPs are known for their function in tip-growing cells like pollen tube and root hair cells as well as interdigitating epidermal pavement cells by regulating Actin-MT dynamics [29,30]. Conversion from the inactive GDP- to active GTP-bound form of ROPs is triggered by ROP-GEFs [31]. Additionally, two IQ67 domain (*IQD*) family genes *IQD6* and *IQD18* were also found downregulated in the *RPS5A>>bdl* background. IQD proteins interact with Calmodulin (CaM) signalling modules and are proposed to mediate Ca2+-dependent regulation of MT organization and dynamics [32,33]. IQD proteins are also emerging as key components in ROP signaling by regulating plasma membrane-MT dynamics for localized growth alterations [34].

**Figure 4:**
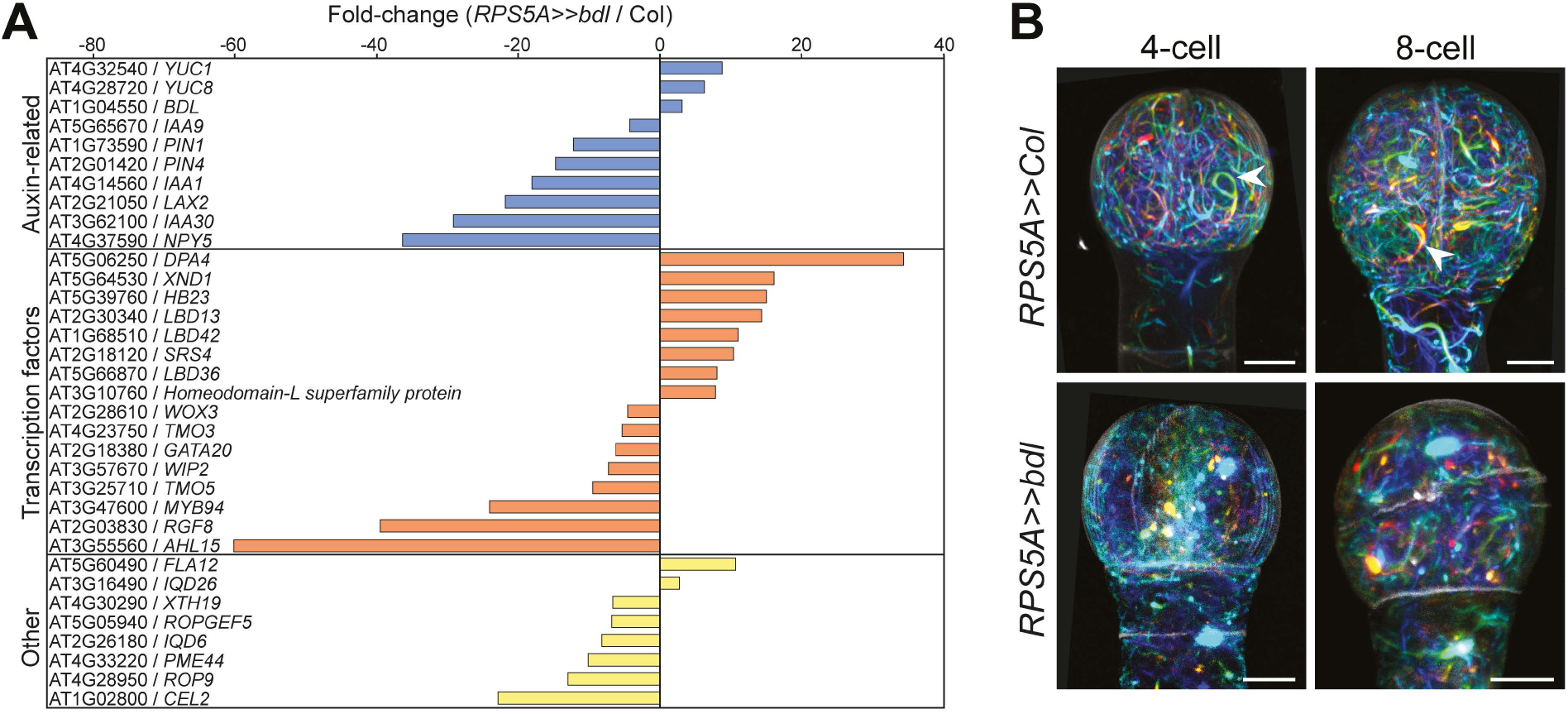
Transcriptome analysis of *RPS5A>>>bdl* embryos. (A) Selected misregulated genes in *bdl*-mutant (*RPS5A>>bdl*) 8-cell embryos. Fold-change values are given for expression levels of genes in the *bdl*-mutant relative to wild-type (*RPS5A>>*Col). (B) Maximum projections of depth colour-coded F-actin stacks visualized using Lifeact-tdTomato (ACE-W14) reporter in wild-type (*RPS5A>>*Col) and *RPS5A>>bdl* embryos. Scale bars: 5 μm

### Auxin response controls cytoskeleton topology in the embryo

The altered expression of a set of genes encoding regulators of Actin and MT cytoskeleton function in auxin-insensitive *RPS5A>>bdl* embryos suggests that auxin response controls these two cytoskeletal structures. We have previously demonstrated that length and degree of MT polymerization is reduced in *RPS5A>>bdl* embryos, and modelling suggested this to contribute to choice of division plane [28]. To address if Actin topology is also altered, we introduced a *pWOX2::LifeAct-tdTomato* (ACE-W14) marker into the *RPS5A>>bdl*background. Previously, we reported thick F-actin bundles in early embryonic cells, which form arches around the nucleus (Fig 4B) [26]. These thick Actin bundles were absent in *RPS5A>>bdl* cells, and in addition, we observed depolymerization defects and loss of dense F-actin meshwork in mutant cells (Fig 4B). Thus, in addition to the effects on the MT cytoskeleton, impaired auxin response causes a disruption of the Actin cytoskeleton in the embryo. By inference, auxin controls the topology of both cytoskeletal structures.

### MT-associated IQD6 mediates auxin action in division control

It is likely that the influence on cytoskeleton function that auxin exerts is mediated by the genes identified as being downregulated in *RPS5A>>bdl* embryos. Here we focused on the *IQD6* gene, which was strongly downregulated (Fig 4A). Previously, inhibition of auxin response on other developmental contexts had been shown to affect the expression of several *IQD* family members [35–38]. Indeed, apart from *IQD6* and *IQDĩ8*, we also observed downregulation of several *IQD* family genes in the *RPS5A>>bdl-mutant* background (Fig S2). We first determined the subcellular localization of IQD6 protein, as well as its close homologs IQD7 and IQD8, by generating C-terminal fusions with YFP (*pIQDX::IQDX-sYFP*). All three proteins show broad accumulation in embryos and roots, with IQD6 and −7 showing a slight enrichment in the root vasculature (Fig 5A; FigS3 and S4). All three IQD proteins exhibited filamentous localization near cell membranes in early embryonic stages, strongly resembling the cortical MT localization reported for embryos previously. IQD6/7/8 proteins localized to the mitotic spindle, phragmoplast and preprophase band (PPB) (Fig 5B and Fig S4), which was not observed for previously reported IQD protein subclades [37], suggesting a possibility of different function for the IQD6-8 family subclade.

**Figure 5:**
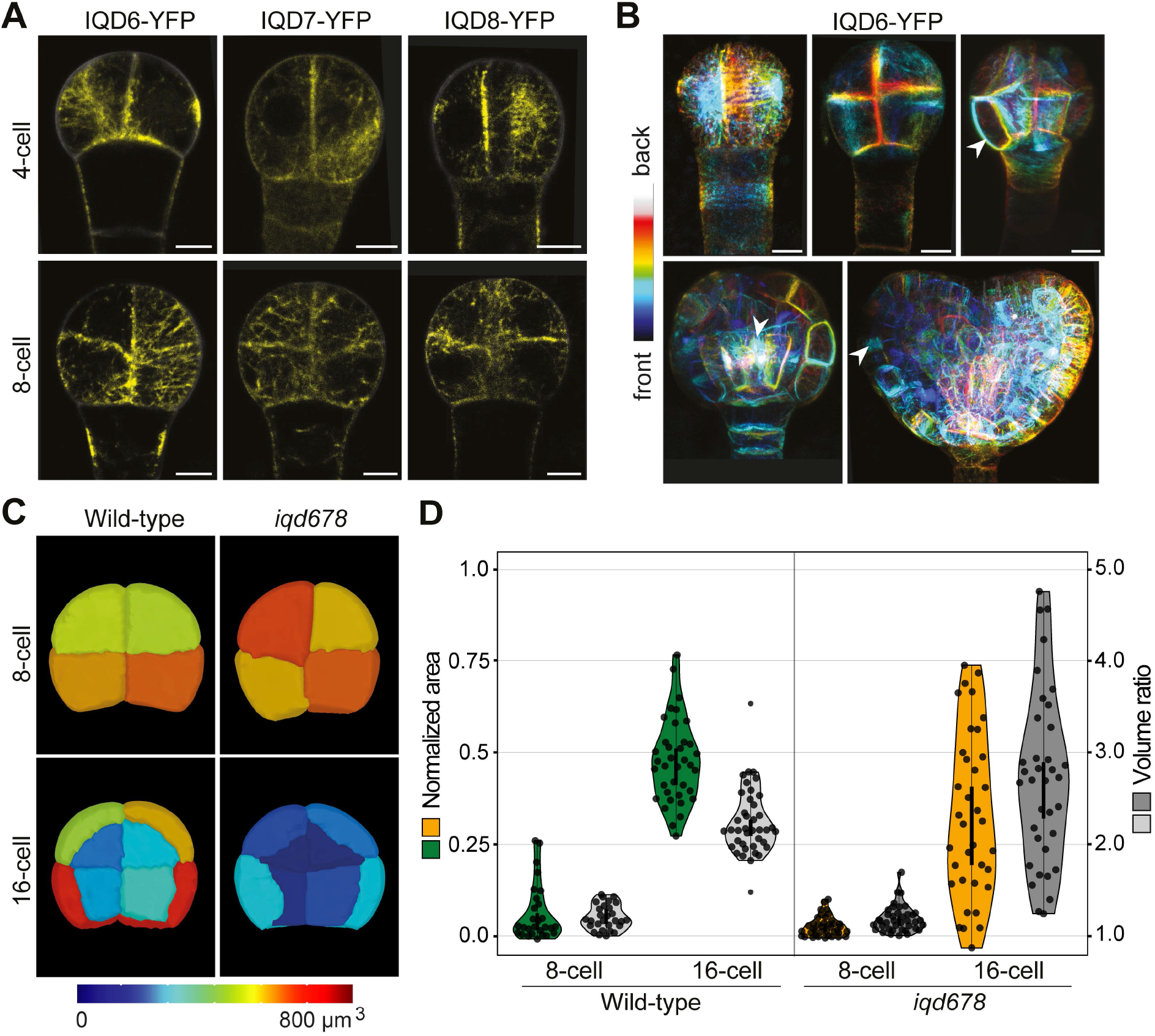
IQD6 mediates auxin response in embryonic division plane control. (A) IQD6, - 7, and −8 visualized in embryos using *pIQDX::IQDX-sYFP* reporter lines show strand-like structures resembling microtubules. (B) Depth colour-coded IQD6-sYFP stacks show localization of protein in 3D. Arrowheads indicate observed localization in pre-prophase bands (PPB), spindles and phragmoplasts. (C) 3D embryo phenotype of *iqd678* mutant embryos with volumetric measurements. Mesh colour per cell corresponds to cellular volume (in μm^3^) indicated in the colour scale. (D) Violin plots representing distribution of division plane areas as a fraction of the smallest (0 on the left y-axis) and largest (1 on the left y-axis) division wall area through the center of the merged volume of two sister cells. Wild-type values are shown in green, *iqd678*-mutant values are shown in yellow. The cell volume ratios resulting from these divisions are represented in grey (light to dark), and values are on the right y-axis. At least 5 individual embryos were used per condition.

To investigate the involvement of IQD6, −7 and −8 in division plane orientation, we analysed the embryos of *iqd678* triple mutants. In 35% of the analysed embryos (Fig S5A), the mutant shows a shift in division plane orientation during different stages, which varies from subtle to more severe defects (Fig S5). The divisions leading to 8-cell embryos are symmetric and use the minimal surface area, similar to wild-type embryos (Fig 5C,D). In contrast, the divisions leading to 16-cell embryos show a high degree of division plane area variation with values spanning across and beyond the normalized areas of wild-type 8-cell and 16-cell stages. (Fig 5C,D). Consequently, volume distribution ratios are also highly variable, from wild-type-like asymmetric divisions to highly symmetric divisions with volume ratios smaller than *RPS5A>>bdl-mutant* embryos (Fig 1B; Fig 5D). These results suggest that IQD6-8 proteins are involved in cell division placement. However, the variability in division plane parameters indicate that this function is not absolute and may signify the involvement of other IQD proteins or additional components. Our analysis also revealed that even when cell division planes and volumes are heavily skewed, the divisions leading to 16-cell stage can still be asymmetric, generating protoderm and inner cell layers (Fig 5C). Regardless, MT-binding IQD proteins act downstream of auxin response in controlling cell division orientation in the Arabidopsis embryo.

### Both MT and Actin cytoskeletons contribute to regulated division plane orientation

Disruption of MT and Actin cytoskeletons is correlated with altered cell division planes in *RPS5A>>bdl* embryos, and this is in turn coupled to altered expression of genes encoding cytoskeletal regulators. It is thus likely, but not proven, that cytoskeleton topology contributes causally to division plane choice in embryo cells. If this were the case, one would expect direct interference with either cytoskeleton to alter cell division planes. Prolonged treatments with cytoskeleton-destabilizing or -stabilizing drugs in embryos is not trivial, and requires *in vitro* culturing of seeds, that in itself can cause abnormalities [39]. We therefore made use of genetic tools to depolymerize MT or F-actin filaments by expression of the PHS1Δ (MT; [40]) or SpvB (Actin; [41]) proteins. Expression of these proteins was previously shown to be equivalent to treatment with MT- or Actin-depolymerizing drugs and disrupts asymmetric radial expansion and polar migration of nuclei in lateral root founder cells [14]. We generated fluorescently tagged versions of PHS1Δ (mNeonGreen-PHS1Δ) and SpvB (SpvB- mNeonGreen) and used GAL4/*UAS* two-component gene expression to drive their expression in embryos only after fertilization, driven by the *RPS5A* promoter. Expression of both proteins in embryos caused frequent changes in division planes. MT depolymerization through PHS1Δ expression led to defects in 95% of embryos (n=122) and caused oblique divisions (Fig 6A; Fig S6) that superficially resemble those induced by inhibition of auxin response. Depolymerizing Actin through SpvB expression led to essentially indistinguishable defects in 85% of embryos (n=165). Also, here, altered division planes are similar to those observed in *RPS5A>>bdl* and *tir1afb12345* embryos (Fig 6A; Fig S6). Defects were obvious at all stages analysed (Fig 6A; Fig S6). To address if these altered divisions are consistent with a switch from asymmetric to symmetric division, we segmented cells and determined both volume ratio and division wall area. In some embryos, the divisions leading to 8-cell embryos show slightly oblique division planes (Fig 6A; Fig S6 and S7). The division plane was not positioned at the centre of the cell in some cells, leading to daughter cells with variable volume distribution. However, while wall area and cell volume ratio were more variable than in wild-type at 8-cell stage (Fig 6B), there was no consistent switch to altered division plane. At 16-cell stage, the asymmetry and volume distribution among cells were also more variable than in wild-type (Fig 6B). In a small number of cells, division wall surface area was larger than in wild-type, correlating with a small population of cells with more asymmetric division (Fig. 6B). At the same time, a larger fraction of cells had smaller division wall surface area, leading to a lower average value in both *RPS5A>>PHS1Δ* and *RPS5A>>SpvB* embryos (Fig. 6B). Consequently, division asymmetry was also reduced in both transgenic genotypes. Thus, while depolymerization of both cytoskeletons expectedly caused more pleiotropic defects in cell division plane orientation, we observe that these defects include the switch to smaller division wall surface area and loss of asymmetric division. We therefore conclude that regulation of the MT and Actin cytoskeleton is critical for asymmetric cell division in the early Arabidopsis embryo, and that auxin response may indeed regulate division orientation through its effects on the cytoskeleton.

**Figure 6.**
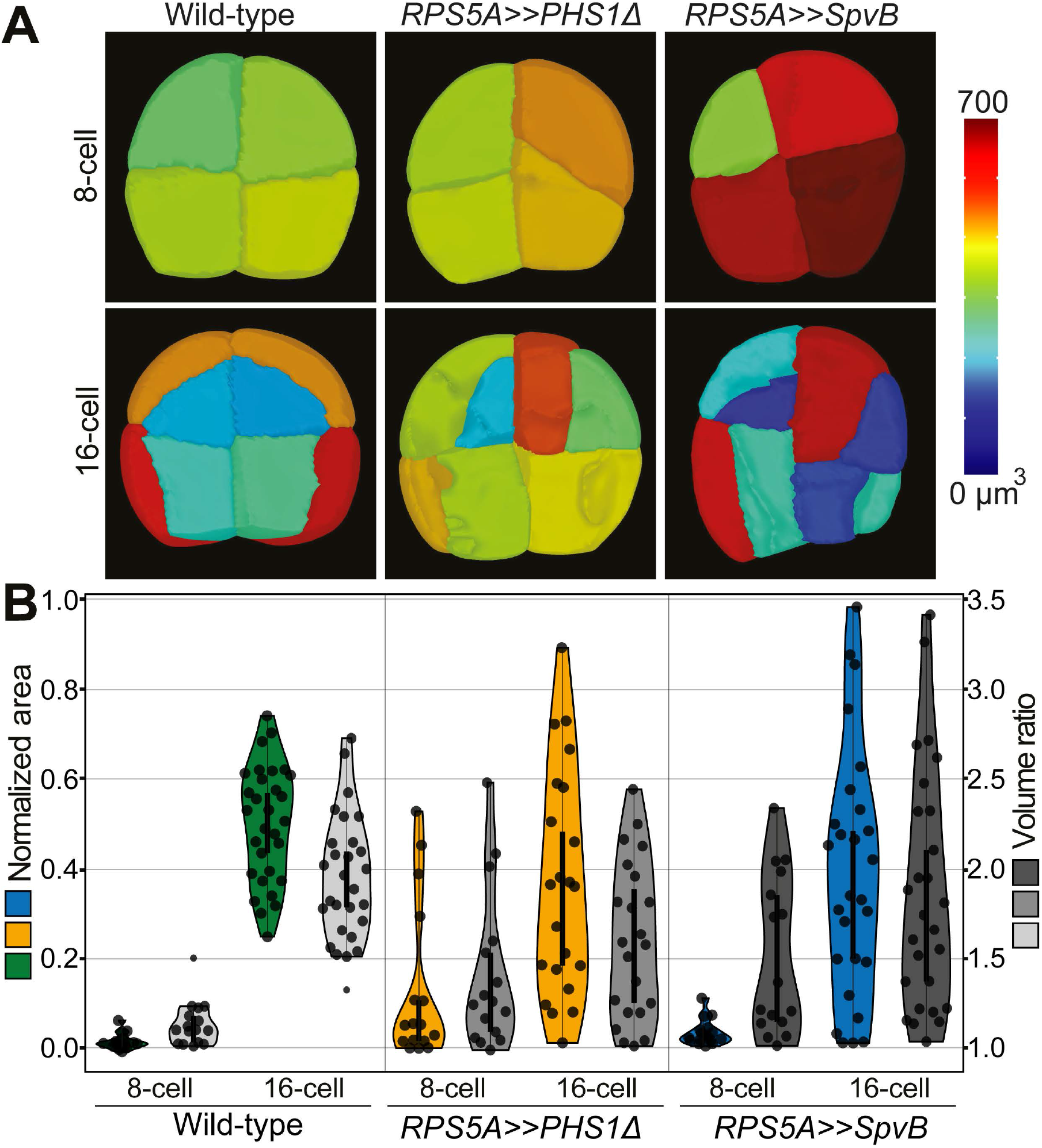
Genetic perturbation of MT and Actin cytoskeleton inhibits asymmetric embryonic divisions. (A) 3D comparison of wild-type, *RPS5A>> PHS1Δ* and *RPS5A>>SpvB-* mutant embryos. Mesh colour per cell corresponds to cellular volume indicated in the colour scale. (B) Violin plots representing distribution of division plane areas as a fraction of the smallest (0 on the left y-axis) and largest (1 on the left y-axis) division wall area through the center of the merged volume of two sister cells. Wild-type values are shown in green, MT- mutant (*RPS5A>> PHS1Δ*) values are shown in yellow, Actin-mutant (*RPS5A>>SpvB*) values are shown in blue. The cell volume ratios resulting from these divisions are represented in grey (light to dark), and values are on the right y-axis. Individual values are shown in the violin plots. At least 4 individual embryos were used per condition.

## Discussion

Incorrect orientation of division plane disrupts tissue patterning and can be deleterious to plant survival. With its highly predictable division pattern, the early Arabidopsis embryo offers an attractive model to study mechanisms underlying oriented division. In this study, we have used the auxin-dependent switch in cell division orientation at the 8-cell embryo stage as a model to understand the control of division placement.

Combining embryo-specific fluorescent cellular reporters with 3D imaging and cell segmentation, we first analysed the role of early polarity, nuclear position, cell shape and auxin mediated cytoskeleton dynamics in orienting the division plane in early embryos. Surprisingly, despite nearly invariant division planes, nucleus position was found to be variable, even in wild-type. Unless we missed a transient stabilization of nuclear position just prior to mitosis, this finding suggests that nuclear position may neither be predictive nor instructive for positioning the division plane in early embryo cells. In contrast, the association of nuclear position and its migration is very clearly demonstrated in zygote, lateral root founder cells and stomatal lineage [13,14,17,42]. Each of those cells however, represent growing cells with large aspect ratio or volume. Since the molecular mechanisms that connect or correlate nuclear position and division plane selection are not yet known, it is not clear if multiple mechanisms exist, or whether the cases in which nuclear movement is observed simply represent exaggerations of the same mechanism operating in embryo cells.

What is evident though, is that changes in cell shape between wild-type and auxin-insensitive embryos correlate with altered division planes. This identifies the control of cell shape as a mediator of division plane choice. While cell wall biology is complex and multifactorial, a key influence on cell shape is the deposition of cellulose fibers along tracks that are dictated by the CMT filaments. Thus, MT dictate the pattern of cell wall fortification and thereby constrain and bias directional elongation, resulting in cell shape changes [43]. Using markers for MT and Actin, we show that the topology of both is subject to auxindependent regulation. This finding is consistent with the central role for MT and Actin in post- embryonic division orientation control [9,44]. Using a transcriptome profiling strategy on 8-cell wild-type and mutant embryos, we identified auxin-dependent genes with known functions in cytoskeleton reorganization. Characterization of IQD6 and its family members imply a significant role in orienting the division plane. While we did not explore other auxin-dependent genes here, several link to the processes identified to be critical to division orientation control. Firstly, a ROP11-IQD13 signalling module was found important for localized growth changes in the formation of xylem pits by organizing CMTs [34]. ROPs are well known to regulate cytoskeleton dynamics during tip growth and in pavement cell interdigitation [45]. In this context, the essential role of ROPs could be keeping the homeostasis of CMTs during the early embryonic stages. CMT stability and polymerization dynamics regulate the PPB formation [46,47]. Simulation studies of CMTs on segmented embryonic cell shapes identify transient auxin-mediated CMT stabilization as a plausible mechanism in division plane orientation [28]. Thus, ROP-mediated cytoskeleton dynamics may play a critical role in fine tuning of the PPB for asymmetric orientation. Secondly, the identification of a set of cell wall-related enzymes suggests that auxin regulation may also directly control wall biochemistry. Recent work has revealed the significance of pectin and xyloglucan in cell wall integrity and remodelling. Methyl esterification status of pectin determines the plasticity of the cell wall and defects lead to severe phenotypes in post-embryonic tissues [48]. These studies represent the wall reorganization effects at tissue and organ level but our current knowledge about the cell wall remodelling by PME or XTH in confined cellular mechanisms like division plane orientation remains poor. The current study opens new avenues for answering these intriguing questions.

Our analysis revealed that outside-inside cell polarity establishment is independent of transcriptional auxin response. We focused on this axis of polarity since the normal division at 8 cell stage aligns with this axis. Thus, unless the outside-inside polarity system has multiple (auxin dependent and independent) branches, auxin acts to control division downstream of this polarity axis. Since no regulators of this axis with function in the embryo have yet been identified, it is at present unknown how this polarity axis still biases the choice of the division plane, and it will be interesting to see how this interacts with the auxin-dependent control of cytoskeleton dynamics and cell shape. Recently, we identified a family of SOSEKI polarity proteins, of which at least two members are transcriptionally controlled by auxin response [49]. Thus, at least part of the polarity system is dependent on auxin input, Misexpression of the SOSEKI1 protein causes oblique cell divisions, suggesting a link to the division orientation machinery. However, it is equally likely that SOSEKI1 affects the CMT or cell shape, and only indirectly influences division plane. Further investigation of this protein family should help resolve how the different cell polarity systems are linked to division control in the embryo.

## Materials and Methods

### Plant Material and Growth Conditions

Arabidopsis ecotype Columbia-0 (Col-0) was used for generating all the transgenic lines. Plants were grown at a constant temperature of 22 °C with a 16-hr light/8-hr dark cycle. Surface sterilized Arabidopsis seeds were subsequently placed on half-strength Murashige and Skoog (MS) medium with agar. After a 48-hour vernalization and 10 days of growth on plates, seedlings were transferred to soil. *tir1 afb* hexuple mutant seeds [24] were kindly provided by Mark Estelle and Michael Prigge (UCSD). ACE-W03 (*pWOX2::BOR1-mCilrine*). ACE-W11 (*pWOX2::AtNUP54-GFP*) and ACE-W14 (*pWOX2::Lifeact-tdTomato*) were previously described [26]. For all crosses *RPS5A-GAL4 (pRPS5A::GAL4-VP16*) [50] was used as female parent. For *bdl* embryo geometric analysis, F1 seeds of cross between RPS5A-GAL4 and Col- 0 or UAS-bdl [51] were used. For nuclear position, F1 seeds of cross between *pWOX2::AtNUP54-GFP* (RPS5A-GAL4) and Col-0 or UAS-bdl were used. For early polarity analysis, F1 seeds of cross between *pWOX2::BOR1-mCitrine* (RPS5A-GAL4) and Col-0 or UAS-bdl were used. For F-actin topology, F1 seeds of cross between *pWOX2::Lifeact-tdTomato* (RPS5A-GAL4) and Col-0 or UAS-bdl were used. For analysis of CMT F1 seeds of cross between*pUAS::PHS1ΔP-mNeonGreen* and Col-0 or RPS5A-GAL4 were used.

For analysis of F-actin F1 seeds of cross between *pUAS::mNeonGreen-SpvB* and Col-0 or RPS5A-GAL4 were used. Seeds of wild-type (Col-0) and T-DNA insertion lines for IQD6 (At2g26180, *iqd6*: SALK_137365), IQD7 (At1g17480, *iqd7*: SALK_025224) and IQD8 (At1g72670, *iqd8*: SALK_107689) were obtained from the Nottingham Arabidopsis Stock Center. All lines were backcrossed at least once with Col-0 and subsequently *iqd6*, *iqd7*, and *iqd8* were crossed among themselves to generate *iqd678* triple mutant.

### Construction of Vectors and Transformation

Plasmids were cloned based on previously described ligation-independent cloning methods and vectors [52]. Whole genomic IQD-sYFP fusions were prepared by cloning up to 3kb of promoter including downstream genomic region up to the stop codon into the pPLV117, containing a super Yellow Fluorescent Protein (sYFP). To generate *pUAS::PHS1ΔP-mNeonGreen* and *pUAS::mNeonGreen-SpvB* plasmids, PHS1ΔP-mNeonGreen and mNeonGreen-SpvB sequences were made by overlapping PCR and introduced into HpaI linearized pLV32. All oligonucleotides used in this study are listed in primer table S1. All constructs were confirmed by sequencing and transformed into Arabidopsis using floral dipping [53]. IQD-sYFP fusions, *pUAS::PHS1ΔP-mNeonGreen* and *pUAS::mNeonGreen-SpvB* were transformed into the Col-0. ACE plasmids [26] were transformed into homozygous RPS5A- GAL4 (*pRPS5A::GAL4-VP16*) driver line.

### Microscopy and image analysis

Embryos were stained by the modified Pseudo-Schiff propidium iodide (mPS-PI) staining method described in [12] with the following modification: An extra treatment with 1% SDS and 0.2 M NaOH for 10 minutes at 37 °C was added after fixation. The stained ovules/embryos were mounted in a drop of chloral hydrate in a well generated by SecureSeal^™^ round imaging spacers (20mm, Thermofisher) and observed by confocal microscopy taking z-stack images. A series of 2D confocal images were recorded at 0.1 μm intervals using a Leica TCS SP5II confocal laser scanning microscope with a 63 × NA = 1.20 water-immersion objective with pinhole set to 1.0 Airy unit. PI was excited using a diode laser with excitation at 561 nm and detection at 600-700 nm.

Embryo samples were prepared as described in [26]. Images for qualitative purpose were acquired in 8-bit format, images for segmentation were acquired in 16-bit format. Images were acquired using a Leica TCS SP5II confocal laser scanning microscope with 63x NA=1.2 water objective with pinhole set to 1.0 Airy unit. mGFP and mCitrine were excited by an Argon-ion laser and tdTomato and SCRI Renaissance Stain 2200 (SR2200) (Renaissance Chemicals, http://www.renchem.co.uk/) were excited using a diode laser, and their emissions were detected sequentially with a Leica HyD in photon counting mode. Excitation and detection of fluorophores were configured as follows: mGFP was excited at 488 nm and detected at 498-528 nm; mCitrine was excited at 515nm and detected at 520-540nm tdTomato was excited at 561 nm and detected at 571-630 nm; Renaissance 2200 was excited at 405 nm and detected at 430-470 nm. Line accumulation was set to 4, 4, and 2 for mGFP, tdTomato, and SR2200, respectively. For qualitative results description of F-actin and nuclear structures, maximum projections were generated. For these stacks, background signal outside of the embryo were subtracted, and remaining embryonic signal was multiplied 2-4 times up until signal saturation. All image processes and measurements were conducted via Fiji.

### 3D cell segmentation and nuclear position measurements

For segmentation, in MophoGraphX (MGX) [25], confocal image stacks (TIF) were Gaussian blurred using sigma value 0.6 μm, subsequently we applied the ITK watershed auto-seeding with level threshold value in the range 300–1500 and default smoothing levels. Segmented bitmap stacks were manually corrected for oversegmentation errors within MGX by fusing together multiple labels into the single cells, which were represented using a combination of the select and paint bucket tools in MGX [54,55]. Then, we approximated the segmented cells by creating triangulated surface meshes using marching cubes 3D with cube size of 1. Nuclear measurements were performed on segmented meshes created using the same segmentation method described above using the nuclei marker channel. Cell and nucleus centroid positions were determined in MGX by calculating the centre of gravity of their triangulated surface meshes. Organ centric directions were determined in the same way as described in the 3D Cell Atlas Add-on for MGX [56] by manually placing a straight line through the embryo using the “Bezier line” in MGX. For each cell then 3 directions relative to this central line were calculated: a longitudinal direction that is identical with the direction defined by the central line, a radial direction that was defined by the cell centroid and its closest point on the central line, and a circumferential direction that was defined by the cross product of the previous two directions. To calculate the distances between cell centroid and nucleus centroid along the longitudinal and radial direction, the scalar product of the vector defined by the centroids and the vector of the respective direction was computed.

### 3D cell morphology measurements

Cell sizes along longitudinal, radial and circumferential directions were computed as described in [56] by shooting rays from the cell centroid along the respective cell directions and their opposites and measure the distance of the two intersection points of the rays with the cellular mesh.

### Shortest division plane estimation and comparison to actual division plane

To compute the relative division plane area, we used the following pipeline in MorphoGraphX which was adapted from [12,55]. First the daughter cells of recently divided cells in segmented meshes were merged. The actual division plane was approximated as a flat wall by computing the principal components of the vertices that were located at the shared border of the two daughter cells. After we simulated a division using this flat wall to determine the surface area of the real division wall (A_real). Then the mean areas of the top 0.1% shortest (A_min) and longest division planes (A_max) in merged cells were determined by sampling of >10000 division directions uniformly spread on the cell volume, going through the center of the actual division wall. Finally, we computed Â = (A_real – A_min) / (A_max – A_min), where Â is the normalized cell wall area, A_min the area of the shortest sampled division planes, A_max the largest sampled division planes, and A_real the area of the flat approximation of the real cell wall.

### Embryo isolation and transcriptome analysis

Ovules were collected from ~60 siliques using vacuum extraction. Siliques were stuck to double-sided tape and sliced open using a needle. Open siliques were submerged in 1x PhosphateBuffered Saline (PBS) buffer and ovules were collected using a vacuum pump through 50 μm filters. Collected ovules were then transferred to Isolation buffer (1x First Strand Buffer (FSB; Invitrogen), 1mM Dithiotreitol (DTT), 4% RNAseLater, MQ), and volume was reduced to ~20 μL. Embryo isolation was performed according to [57] with the following adaptations. A Zeiss Confocor 1 inverted microscope (Carl Zeiss Microscopy GmbH, Jena, Germany) together with an Eppendorf Transferman 4r micromanipulator (Eppendorf AG) and VacuTip II microcapillaries (Eppendorf) were used to isolate about 40-50 washed embryos in 50 μl isolation buffer.

RNA was amplified using the Ovation Pico WTA System V2 (NuGEN, CA, USA), labelled with the ENCORE Biotin Module (NuGEN) and hybridized to Arabidopsis Gene 1.1 ST arrays (Affymetrix, CA, USA) according to the manufacturers protocol. Microarray analysis was performed using the MADMAX pipeline [58] and a custom CDF file (MBNI CustomCDF version 19.0.0) (Dai et al., 2005). Here, all expression values were (quantile) normalised by the Robust multi-array average algorithm (RMA) [59]. Probe sets were redefined using current genome information [60] and re-organized according to TAIR10 gene definitions. Linear models and an intensity-based moderated t statistic approach [61,62] were used to identify differentially expressed genes (probe sets). P-values were corrected for multiple testing using an optimized false discovery rate (FDR) approach [63].

## Acknowledgements

The authors are grateful to Mark Estelle and Michael Prigge (UCSD) for sharing the *tir1 afb* hexuple mutant, to Valentijn Jansen, Joakim Palovaara, Jenny Jansen and Mark Boekschoten for help and support with initial transcriptome analysis and data analysis, to Jos Wendrich for advice and discussions on IQD proteins and to Alexis Maizel for discussions regarding the use of PHS1Δ and SpvB. This work was funded by a fellowship from the German funding agency (DFG; VA 1156/1-1) to P.V., and grants from the Netherlands Organization for Scientific Research (NWO; ALW Open Competition grant 824.14.009) and the European Research Council (Starting Grant “CELLPATTERN”, contract number 281573; Advanced Grant “DIRNDL”, contract number 833867) to D.W.

## Author contributions

P.V.: Conceptualization, Resources, Investigation, Visualization, Formal analysis, Funding acquisition, Writing – Original Draft; T.d.Z.: Conceptualization, Resources, Investigation, Visualization; Formal analysis; S.S.: Methodology, Software, Formal analysis; K.B.: Resources; C.-Y.L.: Investigation; R.S.: Software, Supervision; D.W.: Conceptualization, Supervision, Funding acquisition; All authors: Writing – Review & Editing.

## Supplementary material

This manuscript contains 7 Supplementary Figures, one Supplementary Table and one Supplementary Data file. The transcriptome data have been deposited in the NCBI Gene Expression Omnibus, and are accessible through accession number GSE165986.

---

**Figure S1:**
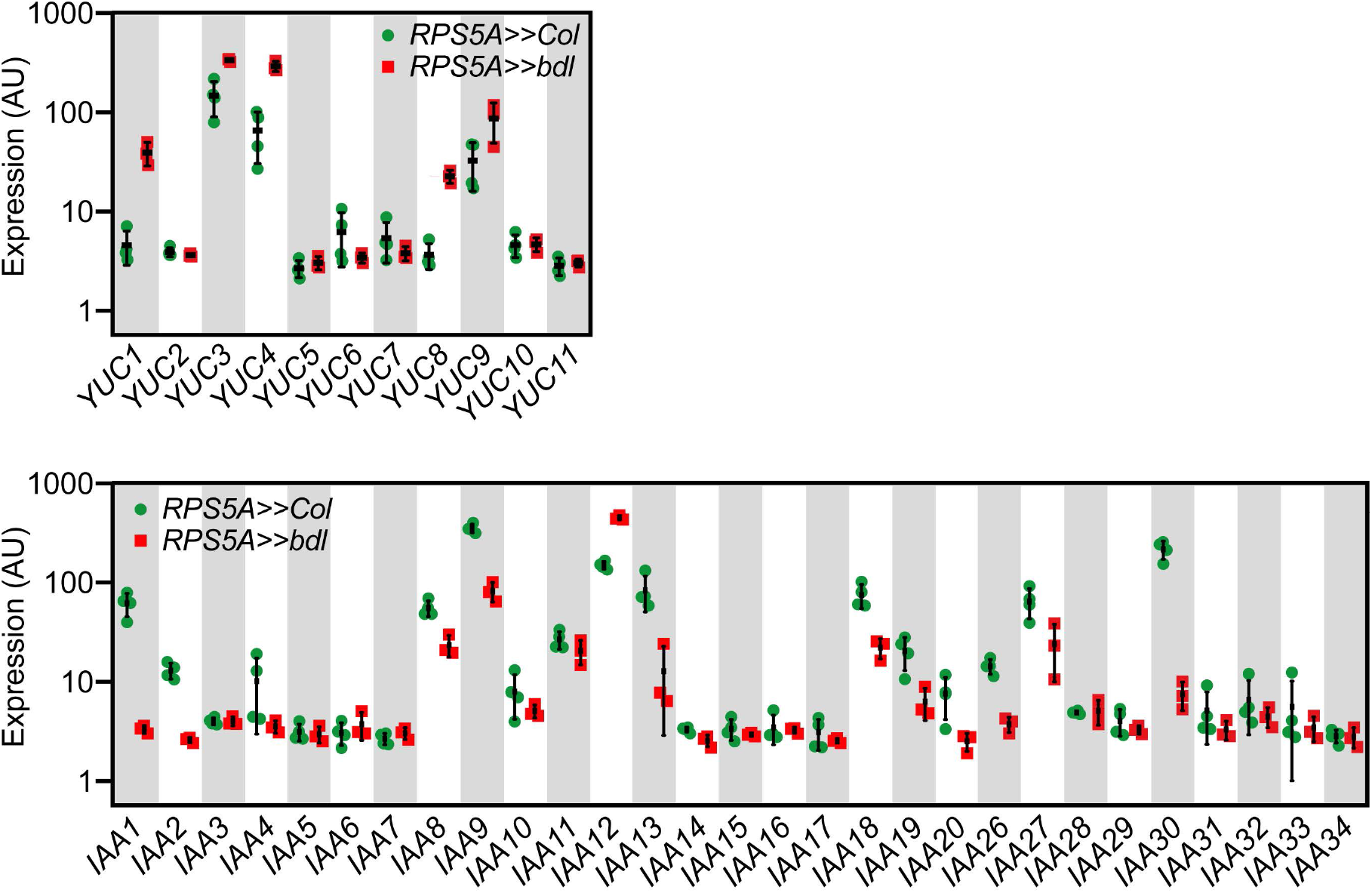
Differential expression of *YUC* and *AUX/IAA* genes in *RPSS5A>>bdl*mutant embryos. Note that 5 out of 11 *YUC* genes (*YUC1,3,4,8,9*) are upregulated in *RPS5A>>bdl* background compared to *RPS5A>>Col* control embryos. Many of the *Aux/IAAs* are downregulated (*IAA1,2,4,8,9,13,18,19,20,26,27,30*) in *RPS5A>>bdl* embryo*s* except *BDL (IAA12*), which is highly upregulated. Those Aux/IAA’s whose expression is not altered have very low absolute expression levels, and can essentially be considered “not expressed” in the embryo.

**Figure S2:**
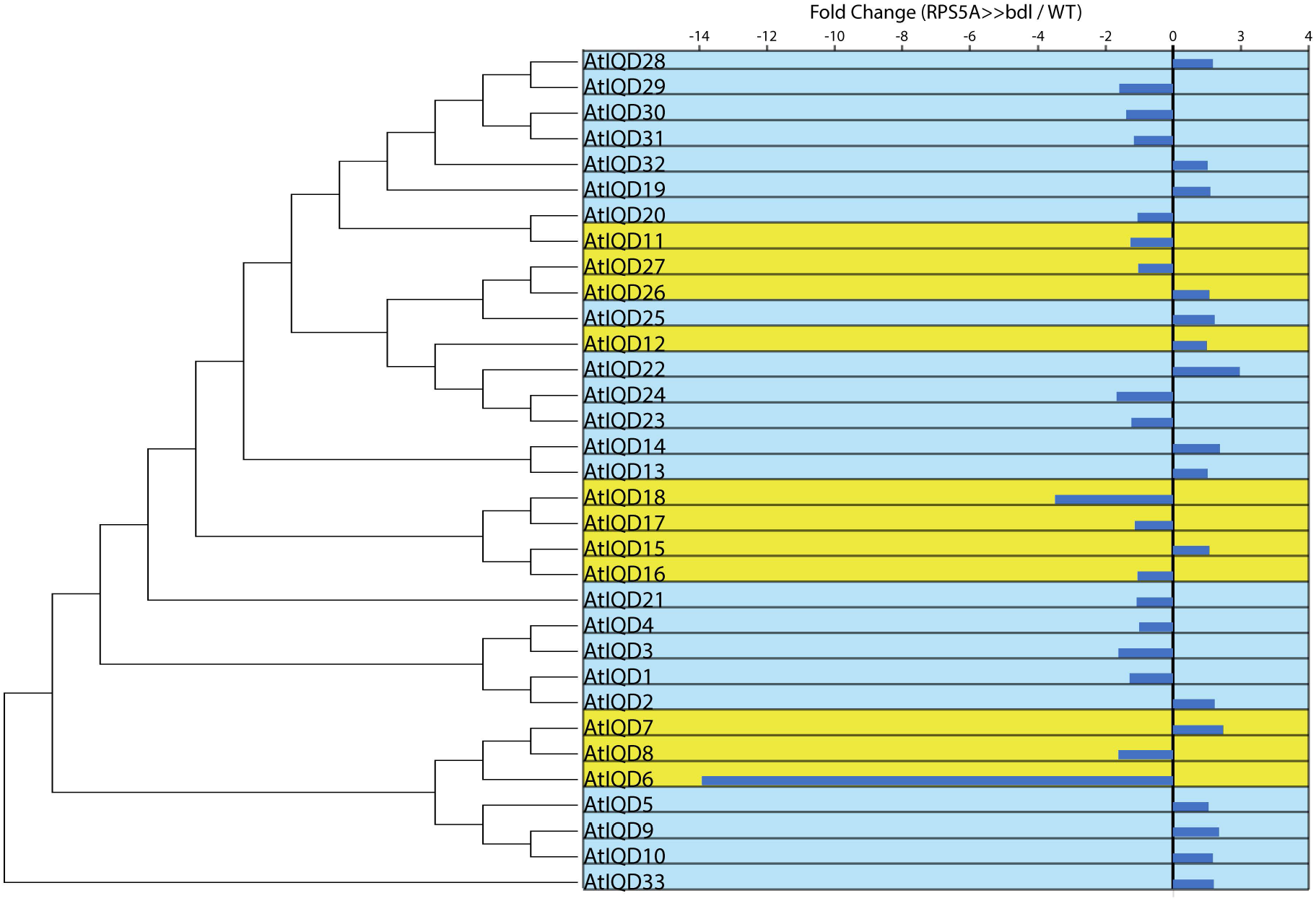
Phylogenetic tree of all Arabidopsis *IQD* genes, rooted to *IQD33*, combined with their misexpression in 8-cell *bdl (RPS5A>>bdl*) embryos. Fold-change values are given for expression levels of genes in *RPS5A>>bdl* mutant embryos, relative to wild-type (*RPS5A>>col*). Yellow boxes indicate subclades previously shown to be misregulated in auxin-related datasets [35—38].

**Figure S3:**
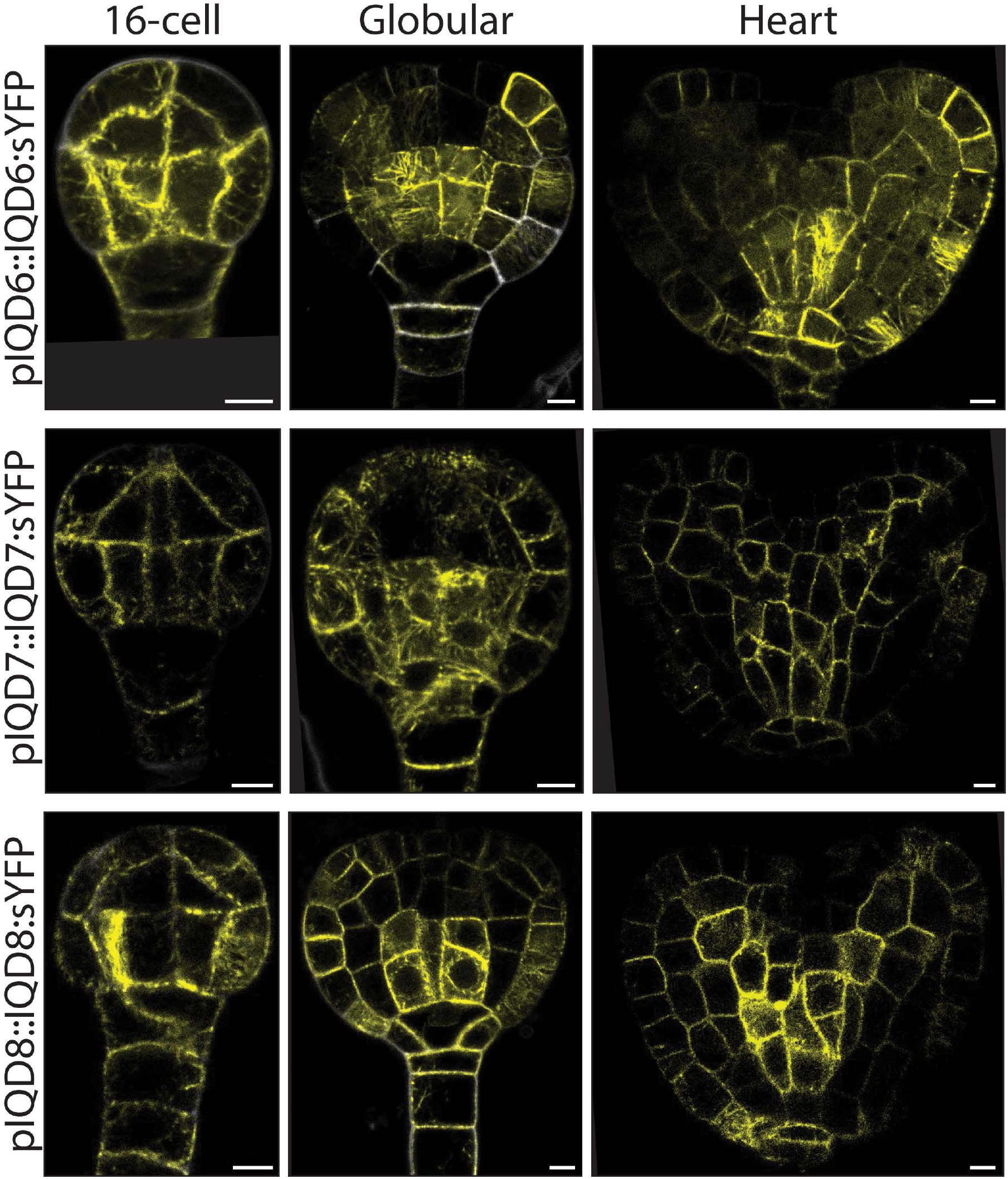
IQD subcellular protein localization during Arabidopsis embryogenesis. IQD6, −7, and −8 are visualized using *pIQDX::IQDX-sYFP* reporter lines. Scale bars: 5 μm.

**Figure S4:**
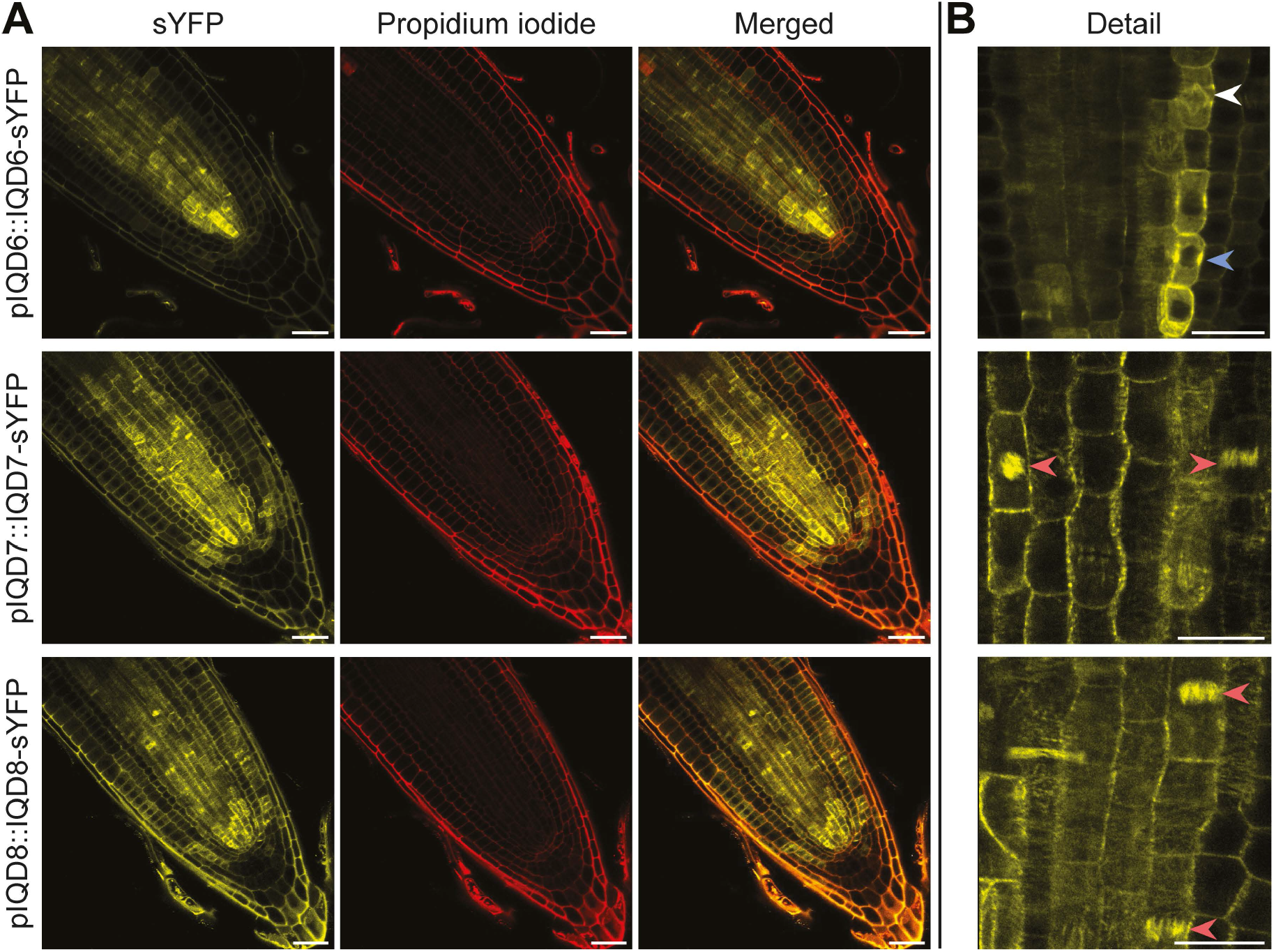
IQD subcellular protein localization in 5-day old Arabidopsis roots. (A) IQD6-, 7-, and 8 are visualized using a *pIQDX::IQDX-sYFP* reporter line merged with membrane visualisation using Propidium Iodide (PI) staining. Scale bars: 30 μm. (B) Details of roots with arrowheads indicate observed localization of protein in preprophase bands (PPB; blue), spindles (white) and phragmoplasts (red). Scale bars: 15 μm

**Figure S5:**
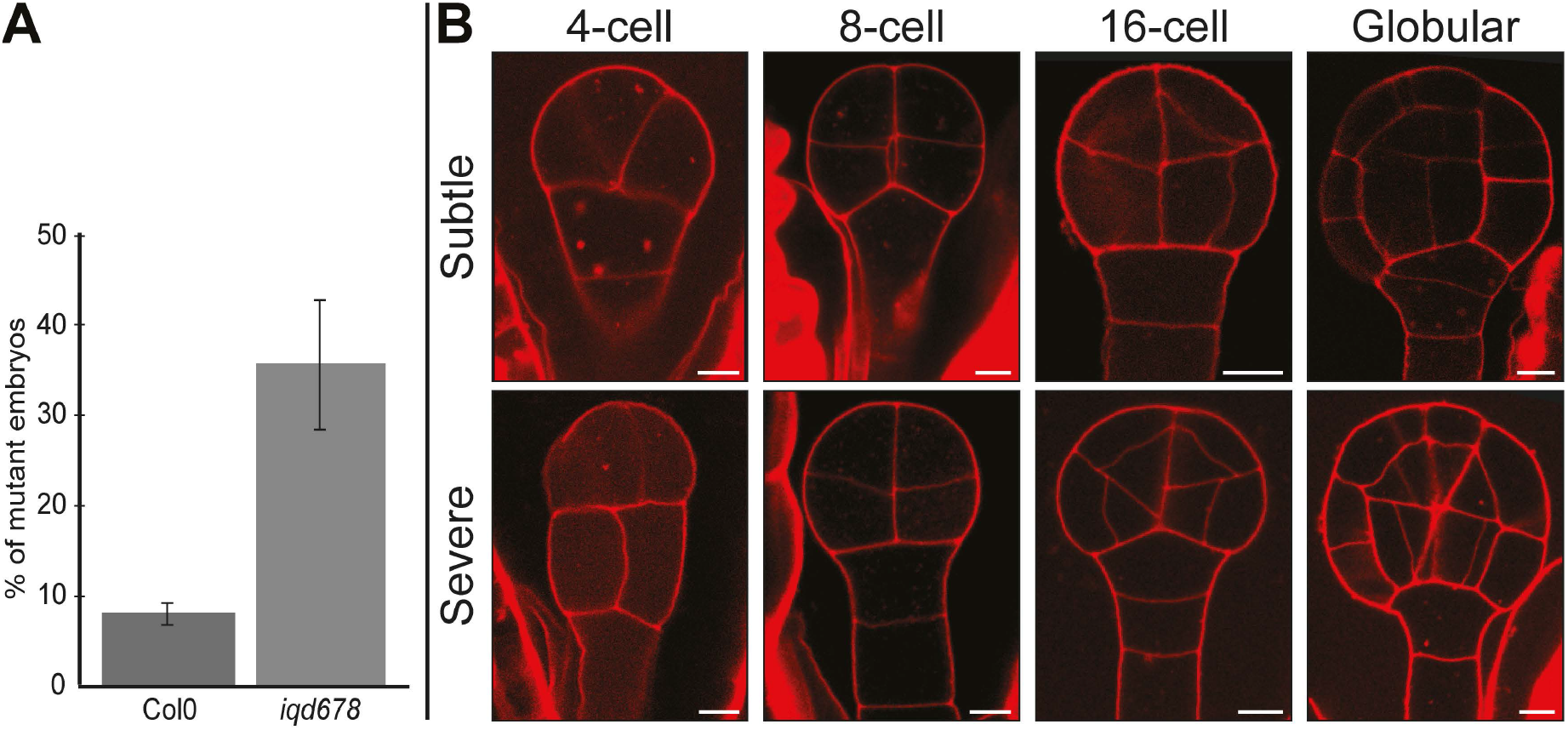
2D embryo phenotype for the *iqd678* mutant. (A) Quantification of skewed division planes in early embryos. Quantification is based on visual inspection of at least 250 individual chloral hydrate-cleared embryos. (B) Embryos can show either a subtle or severe phenotype (Stained using modified Pseudo-Schiff propidium iodide (mPS-PI) method). Scale bars: 5 μm.

**Figure S6:**
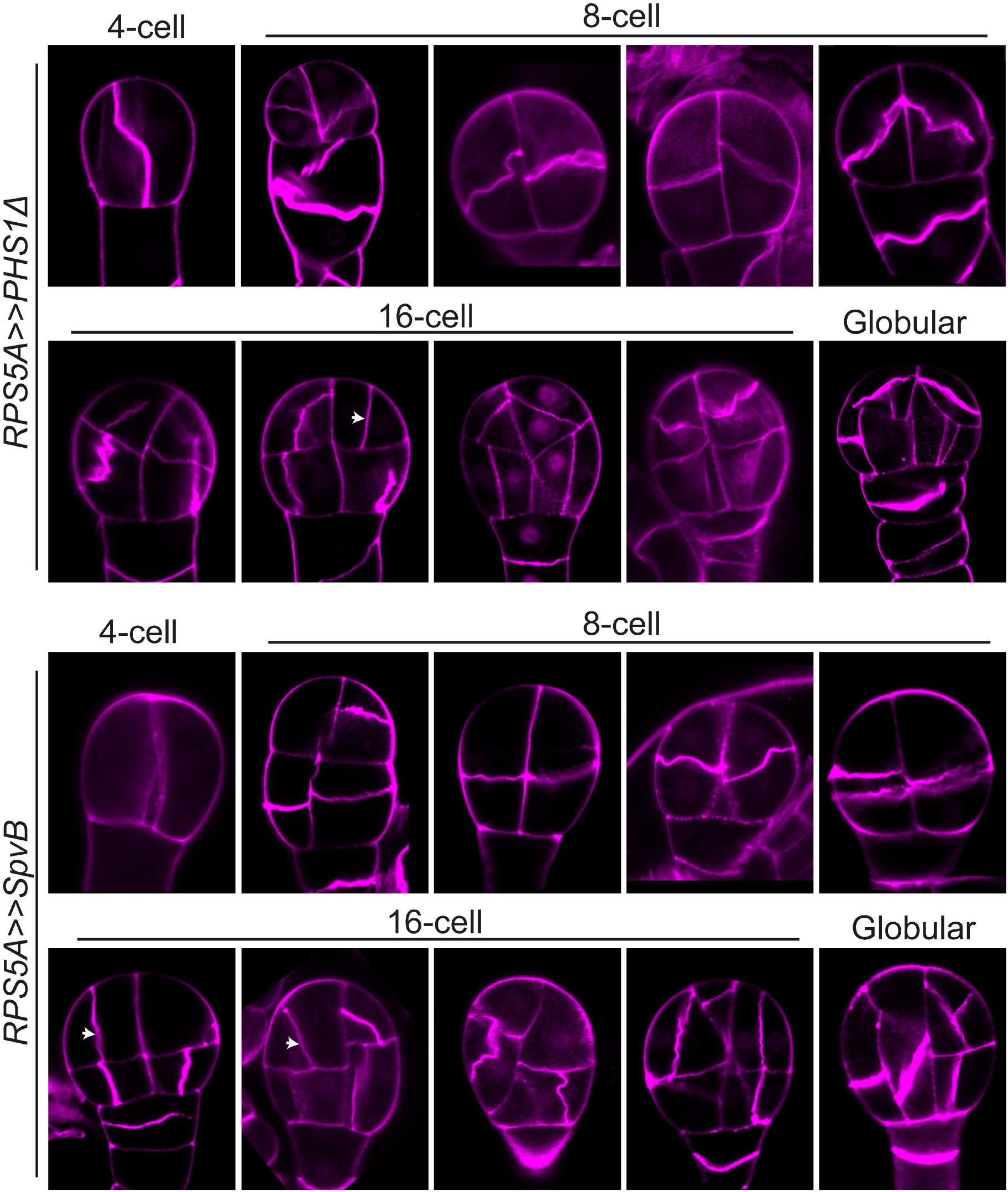
2D embryo phenotype for *RPS5A>>PHS1Δ* and *RPS5A>>SpvB-* mutant embryos. Embryos exhibit high variation of division plane defects among samples. Arrows indicate *bdl-*like division plane defects. Stained with SCRI Renaissance Stain 2200 (SR2200).

**Figure S7:**
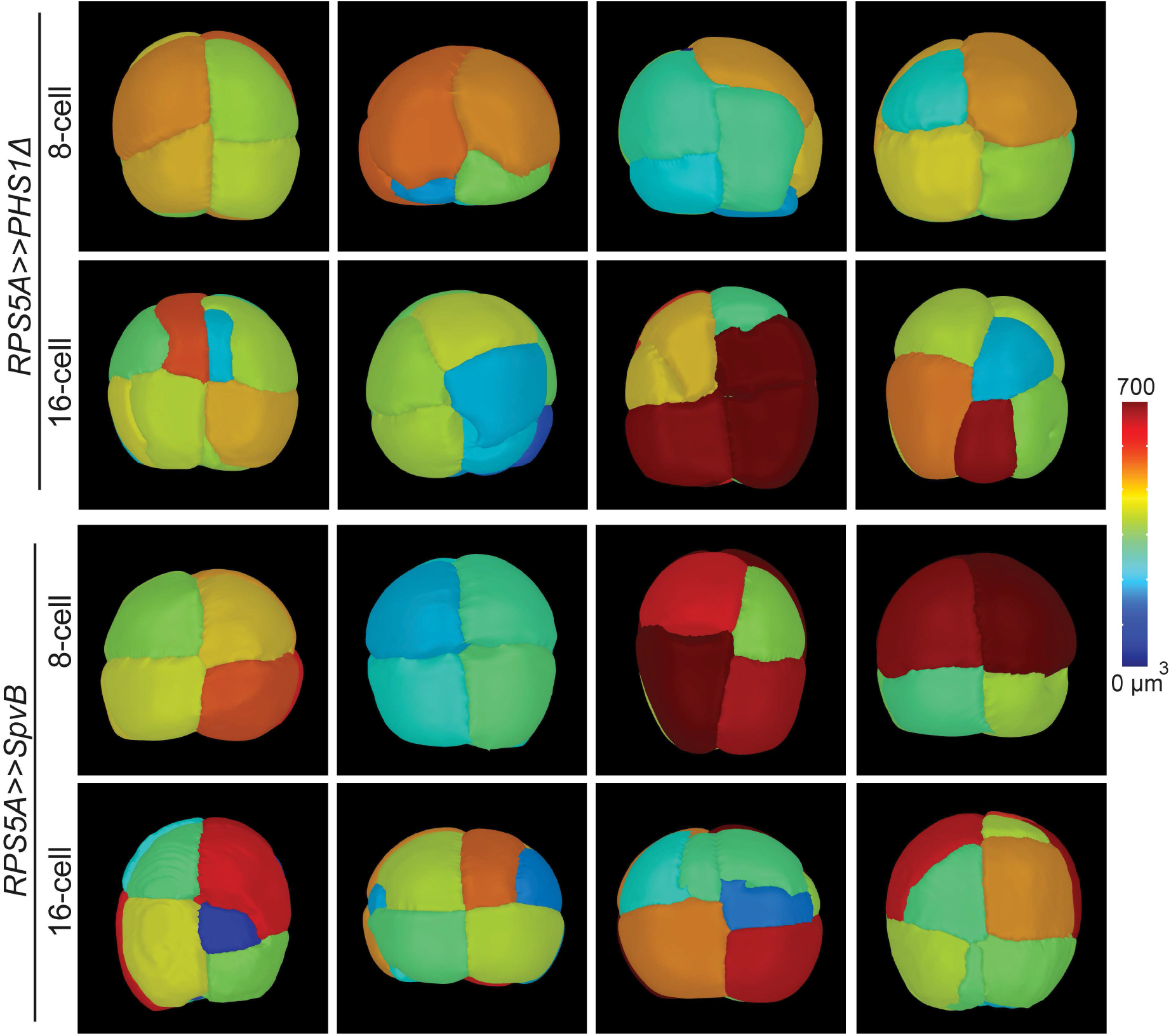
3D phenotype of *RPS5A>>PHS1Δ* and *RPS5A>>SpvB-* mutant embryos with volumetric measurements. Mesh colour per cell corresponds to cellular volume (in μm^3^) indicated in the colour scale. Samples used for geometric analysis in Fig 4B.

**Primer Table S1:**
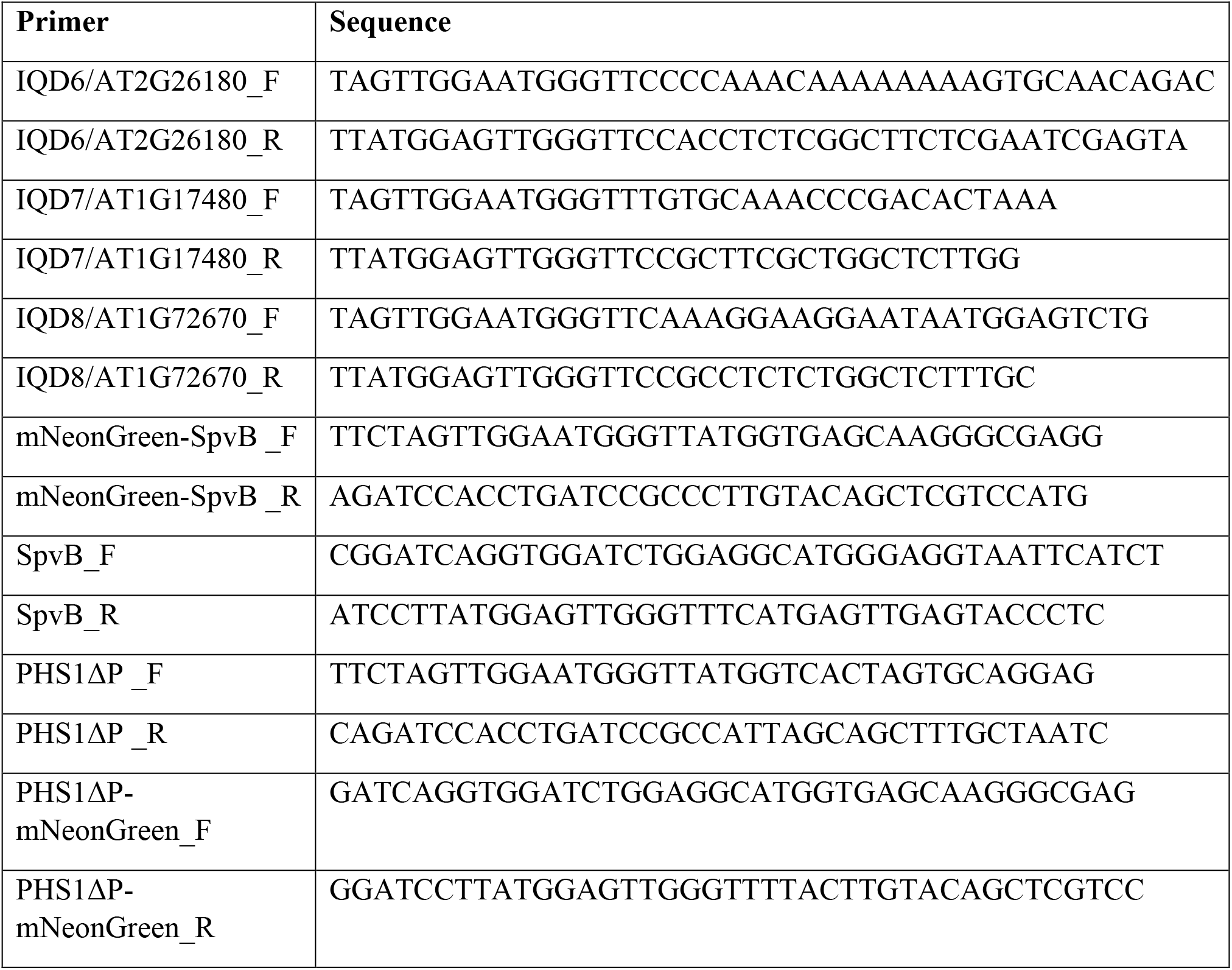

